# Statistical Association Mapping of Population-Structured Genetic Data

**DOI:** 10.1101/069658

**Authors:** A. Najafi, S. Janghorbani, S. A. Motahari, E. Fatemizadeh

## Abstract

Association mapping of genetic diseases has attracted extensive research interest during the recent years. However, most of the methodologies introduced so far suffer from spurious inference of the disease-causing sites due to population inhomogeneities. In this paper, we introduce a statistical framework to compensate for this shortcoming by equipping the current methodologies with a state-of-the-art clustering algorithm being widely used in population genetics applications. The proposed framework jointly infers the disease causal factors and the hidden population structures. In this regard, a Markov Chain-Monte Carlo (MCMC) procedure has been employed to assess the posterior probability distribution of the model parameters. We have implemented our proposed framework on a software package whose performance is extensively evaluated on a number of synthetic datasets, and compared to some of the well-known existing methods such as STRUCTURE. It has been shown that in extreme scenarios, up to 10 – 15% of improvement in the inference accuracy is achieved with a moderate increase in computational complexity.

## 1 INTRODUCTION

LARGE-SCALE projects in life sciences, such as Human Genome Project [1] and HapMap project [2], [3], have provided biologists and computer scientists with an invaluable foundation for study and research. In addition, emergence of high throughput sequencing technologies has paved the way to solve the main problems in biology, such as Genome-Wide Association Study (GWAS) [4]. The basic purpose of GWAS is to infer statistical associations between different regions of genome and specific physical or behavioral phenotypes present in living organisms. In many medical applications, as of those considered in this paper, the aforementioned phenotypes are the affection by or vulnerability to a particular genetically-initiated disease. In other words, the goal of GWAS would be to assign specific sites in the DNA sequence, Single Nucleotide Polymorphism (SNP) data or even intensity levels of a microarray experiment to the causal factors underlying a specific disease [5]. During the recent years GWAS methods have been successful in identifying many causal factors for different types of diseases. However, despite major advantages, traditional methods in this area suffer form critical drawbacks.

First, most traditional GWAS frameworks consider genetic variants, such as SNPs, separately and neglect the effect of their biochemical dependencies, a phenomenon called *epistasis* [6]. This premise may lead to spurious results in occasions where multiple loci are involved in the formation of a complex disease. In other words, multigenetic factors exist in many complex abnormalities since multiple pathways may control a specific biological reaction. In this regard, alternation of each pathway may result into the same disease with highly similar symptoms. This shortcoming usually increases the false discovery rate in limited sample sizes. Recently, a number of researchers have set out to alleviate this problem by introducing various statistical and/or experimental tools[7], [8].

The second major shortcoming, which has triggered the idea behind the current paper, is the assumption of genetic homogeneity for the population under study. This assumption is not plausible in real-world datasets since different individuals may have come from different ancestral origins. In such scenarios, also known as “cryptic populations”[9], attempting a naive association mapping may lead to incorrect outcomes since averaging the statistical results over the whole population produces noisy statistics and decreases the significance levels of the causal genes[10], [11], [12]. In addition, self-reported ancestries often do not provide sufficient evidence[13]. In order to rectify this effect several approaches have been proposed, yet each one suffers from its own drawbacks. In particular, majority of previous algorithms use a population stratification strategy to cancel the effect of data structures by clustering the individuals first, and then feeding each cluster to a GWAS module, separately [14]. However, the unsupervised clustering phase ignores the information provided by the disease labels, and its accuracy will highly depend on allele frequencies. This will degrade the performance of the overall framework over small datasets.

In this paper, we address all of the mentioned problems by proposing a novel method for association mapping in the presence of hidden population structures. In other words, it has been assumed that the population under study consists of numerous latent sub-populations with different genetic backgrounds. More importantly, these differences in genetic ancestries are assumed to correlate with distinct genetic vulnerability to the disease, resulting in different disease infection models for each sub-population. We have shown that integration of the distinctions both in allele frequencies and also the disease models highly improves the identification of latent structures as well as causal genetic factors. We have developed a model-based statistical framework which combines genotype clustering algorithms with current association mapping strategies to form a unified mathematical tool with a significantly higher accuracy.

The paper is organized as follows. In Section 2 related works in association mapping and GWAS are reviewed. Section 3 explains the basic ideas and mathematical notations in this work. In Section 4, the proposed model is explained, while in Section 5 the statistical inference of model parameters from data is discussed. Section 6 presents our computer simulations and experimental results. Conclusions are made in Section 7.

## 2 BACKGROUND AND RELATED WORKS

So far, GWAS methods have been conducted on a wide range of abnormalities and resulted in numerous scientific discoveries. For instance, in[15], [16] and [17] a number of causal loci for Type I diabetes have been identified, while in[18], [19], [20], [21] the same is carried out for type II diabetes. GWAS methods have also been put to use for more complicated anomalies such as different types of cancer[22]. Researchers in[23], [24], [25], [26] investigate the causal factors for breast cancer, while[27], [28], [29] have identified a number of disease-causing genes for prostate cancer. The application of GWAS methods are extended to genetically initiated mental disorders as well, such as Parkinson’s disease as in[30], [31], [32], Bipolar Disorder as in[33], [34], and Schizophrenia [35], [36]. Many of such findings are currently being used to diagnose and treat various diseases in gene therapeutic centers worldwide [37].

Despite novel achievements of GWAS methods [38], the effect of population structures may generate spurious results. Based on this motivation, a variety of approaches have been proposed by researchers to solve such problems. One approach is to design family-based studies for association mapping instead of case/control groups. Although several versions of these methods exist[39], most of them are underpowered since the data needed for such methods is difficult to obtain[4], [40].

A class of well-known approaches applies appropriate clustering methods to case/control groups in order to identify the latent structures within data. Such methods are used as a preprocessing stage before the actual association mapping. In particular, principal component analysis (PCA)[41], mixed model approaches[42] and algorithms using the STRUCTURE framework[9] are being widely used. In PCA-based methods, continuous axes of variation with the most amount of information about genetic variability, also known as *principal components* will be determined, which reveal information regarding population structures of the data[41], [43]. A faster and more accurate version is proposed in[44]. More recent studies show that PCA is less robust comparing with nonlinear methods such as spectral dimensionality reduction[45], [46]. Despite of relative improvements in results, top principal components do not necessarily represent true genetic structures since their application lacks an appropriate biological plausibility. The same argument holds for spectral techniques. In fact, they mix structures with long distance LD, family-relatedness or artifacts [47]. In Mixed Model Approaches, the phenotypes are modeled as a mixture of fixed and random effects. These methods, however, may have a lower performance in comparison to their counterparts [47]. Several versions of these methods including [48], [49] or the faster version in [50] have been proposed so far.

Among the most popular approaches is the seminal work introduced in[9] which is known as a state-of-the-art clustering method based on a Bayesian framework, called STRUCTURE. More recent methods motivated by this approach also exist, see[51] and[52]. As suggested in[9], one can apply STRUCTURE to case/control groups in an unsupervised scenario to identify hidden structures. As we will show in this paper, this procedure undermines the true potentials of Bayesian estimation in GWAS methodologies. A combination of the many of the above methods is used in[53] where PCA is combined with Random Forest, and also in[54] where PCA meets Linear Mixed Models.

Our proposed algorithm is built upon STRUCTURE. However, it takes the disease infection labels of a GWAS dataset into account during the clustering phase. This way, the disease infection model, i.e. association mapping, and clustering, i.e. identification of latent population structures, will be carried out simultaneously and interactively.

## 3 PROBLEM FORMULATION

We are interested in finding the causal genomic variants of a particular phenotype in a given population. In most cases of interest, the observable phenotype is the affection by a particular genetic disease. To this end, *N* affected and unaffected individuals are sampled from the population. Each individual is labeled indicating whether or not he/she possesses the phenotype.

Each individual is genotyped at *L* genomic loci. Each locus can take *J* distinct values indicating distinct allele types obtained either from SNP sets or microsatelite data. The data obtained from individuals can be represented by ***D*** = (***X, Y***) where

***X*** ∈ {1, 2, …, *J*}^*N*×*L*^ represents the genotype data of individuals obtained either from the SNP sets or microsatellite data.
***Y*** ∈ {0, 1}^*N*^ demonstrates the the labels showing whether each individual is associated with a particular phenotype, such as a particular disease or not.

As it is mentioned, our primary aim is to infer causal genomic variants of the population given the data *D*. In a simplifying model, all affected individuals share the same set of genomic loci as the cause of the given phenotype. In this case, one can perform several statistical inference strategies to obtain the variants given the data. In our more realistic study, however, people in the population are affected differently due to the fact that individuals are originated from *K* hidden sub-populations and the set of causal variants are different in each sub-population. As discussed before, the presence of such loci in genome is tightly related to genetic evolutionary pathways such as independent genetic drifts. Also, extrinsic evolutionary forces such as natural selection may affect individuals of the same species differently, as a result of the differences in the environmental factors of their habitat.

Our goal is to obtain *K* different sets of causal variants from the data *D*. Note that it is assumed that *K* is known. In practice, the number of sub-population can be inferred via trial-and-error methods.

Sub-populations are differentiated based on their minor allele frequencies (MAF). In other words, associated with each location *j* is a *hidden* number *p_j,ℓ,k_* indicating the frequency of the *ℓ*th allele in the *k*th sub-population. We use an array ***P*** = [*p_j,ℓ,k_*] to indicate the MAF of all subpopulations, i.e.,

> 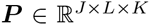 represents the frequency of alleles at each locus for each sub-population.

The *i*th individual is originated from a sub-population *Z_i_* ∈ {1, ⋯, *K*}. We denote the *hidden* vector of associations to sub-populations by ***Z*** = [*z_i_*], i.e.,

> ***Z*** ∈ {1, 2,…, *K* }^N^ represents the sub-population of origin for individuals.

Finally, the model underlying the corresponding complex phenotype for the kth sub-population is denoted by *M_k_*. In particular, *M_k_* indicates the causal genomic loci affecting the kth sub-population. We denote the vector of models by ***M***, i.e.,

> ***M*** = {*M*_1_, *M*_2_,…, *M_K_*} represents the disease-causing model in each sub-population.

In fact, ***M*** models the mathematical relation of disease labels with all other parameters of the problem. In Sections 1 and 2, a concise overview of previously introduced models in GWAS, their cons, pros and computational complexities is presented. The most important assumption in this study, is letting the complex disease model ***M*** to vary for each sub-population. Several recent findings in the pathology of complex genetic diseases support such mathematical assumption[5], [11], [17]. This is due to the fact that functional misbehavior of vital processes in living organisms can occur from multiple sources of genetic abnormalities rather than one.

## 4 THE PROPOSED MODEL

In this section, we provide a probabilistic model governing the main parameters of the problem: ***D*** = (***X, Y***) and ***H*** = (***P,Z,M***). ***D*** stands for data and ***H*** for hidden parameters. Clearly,

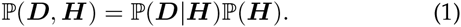

We need to present a model incorporating our knowledge into the priors, i.e., defining 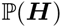, and the way data are generated from the hidden parameters, i.e., defining the conditional distribution 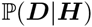.

### 4.1 Modeling of Prior Distributions

We assume statistical independence among prior knowledge of allele frequencies ***P***, information regarding subpopulations of origin ***Z*** and also the disease causing models ***M***, as in [9]. This assumption is biologically plausible since in reality there are not much evidence for statistical linkage of these quantities, i.e.,

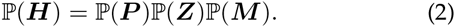

To model 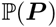, we note that *p_*,ℓ,k_* = {*p_1,ℓ,k_,p_2,ℓ,k_,…, ℓp_J,ℓ,k_* } is a probability distribution and it sums to one. Therefore, similar to [9], we use the Dirichlet distribution to model the allele frequencies:

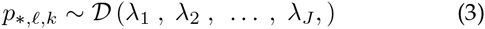

where *λs* are user-specific parameters, all of which can be set to one in case there is no prior information. We also assume that *p_*,ℓ,k_* for *ℓ* and *k* are independent.

Assuming that sub-populations have the same number of individuals, a random individual belongs to the *k*th sub-population with probability 1/*K*. From independent sampling of individuals, hence, we obtain

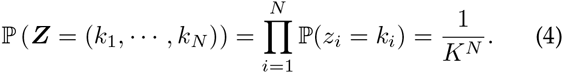

In order to discuss the mathematical models of a complex disease, we first take into account a number of biologically related assumptions. First, it is assumed that all the individuals are labeled in correspondence with one particular genetic disease. Moreover, we assume that the disease of interest has multigene causal pathways. In other words, the biological complexity of the disease strongly suggests that different, and apparently independent genetic abnormalities may lead to the same misfunctionality in body. In this regard, it would be reasonable to assume that different subgroups of a population are associated with different causal factors which justifies decomposing the disease model into *K* independent sub-models, where each submodel is related to a specific genetic sub-population.

Based on the above-mentioned assumptions, a general mathematical model for a complex genetic disease in each sub-population assumes statistical dependence between the disease and a particular group of SNPs or genetic variations. Mathematically speaking, disease-causing sites denoted by 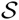 can be decomposed into {*S_1_, S_2_,…, S_k_*}, where *S_k_* includes all the loci associated with the disease in the *k*th sub-population.

Various assumptions regarding |*S_k_*|, i.e. the number of causal loci in the *k*th sub-population, can be made. A naive approach would be to consider single locus hypothesis testing which ignores epistatic relations among genetic sites. More complicated assumptions incorporate investigation of multiple genetic markers instead of one which lead to better results, yet suffer from highly increased computational burdens. We have assumed |*S_k_*| to be drawn from a *Poisson* distribution with an adjustable parameter *η_k_:*

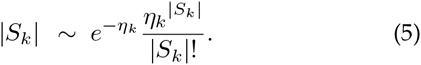

User can choose large values for *η_k_* in order to incorporate more causal variants in the model. This, in turn, increases the computational complexity of the proposed statistical inference schemes.

An appropriate prior for choosing elements in each *S_k_* is to promote those combinations of loci which are physically close to each other in genome. This way local epistatic relations in formation of a complex disease can be appropriately addressed. Therefore, one can rewrite the conditional probability distribution of 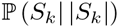 as:

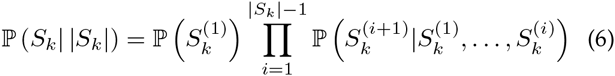

where 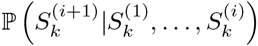 takes non-zero values only for a fraction of 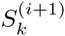 which are located in Δ neighborhood of at least one of the previously determined causal loci, i.e. 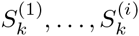. Again, Δ is a user specific parameter and indicates the extent of epistasis in genome which we wish to consider. Based on the above assumptions, the prior distribution for the disease model can be expressed as:

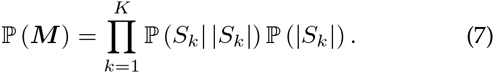

### 4.2 Data Modeling

To model the generation of data given the hidden parameters, we note that

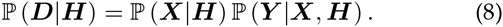

Therefore, we discuss about the two factors separately. First, we argue that

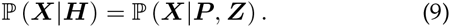

This is due the fact that, in our model, individuals attain their genomic variants from the sub-population that they are originated from, and the disease model only affects people with certain genotypes.

For the sake of simplicity, we assume linkage equilibrium among genetic loci as well as Hardy-Weinberg equilibrium in each sub-population of origin. In addition, the sub-population specific frequencies between different groups are completely independent. In proceeding sections, these assumptions enable us to draw independent samples from the allele frequency distributions. In the absence of linkage disequilibrium (LD), genotype matrix probability distribution can be formulated by a series of multinomial functions as follows:

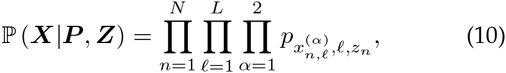

where 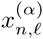 denotes the genotype of the *n*th individual in his/her lth locus of the *α*th chromosome (here we have focused on *diploid* organisms such as humans).

To model the second factor in Equation (8), we note that

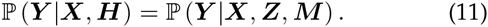

This is due to the fact that whether or not a person is affected is independent of the MAFs of his/her sub-population.

Considering independence in susceptibilities of individuals to the given disease, one can write

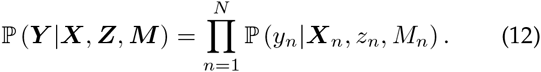

Given *z_n_* and *M_n_*, one can obtain the causal loci of the disease for each person of interest. Let us denote this set of loci for the nth individual by *W_n_*. Obviously, *W_n_* can take *J*^2|*S_z_n__*|^ possible combinations. Each combination for *W_n_* infects the individual with an unknown probability denoted by *F_z_n__* (*W_n_*). Hence,

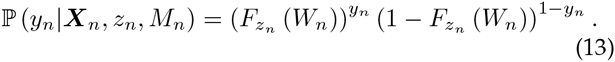

A Bayesian network can be used to capture all the dependencies between data and hidden parameters. Fig. 1 presents the graphical model of this network.

**Fig. 1.**
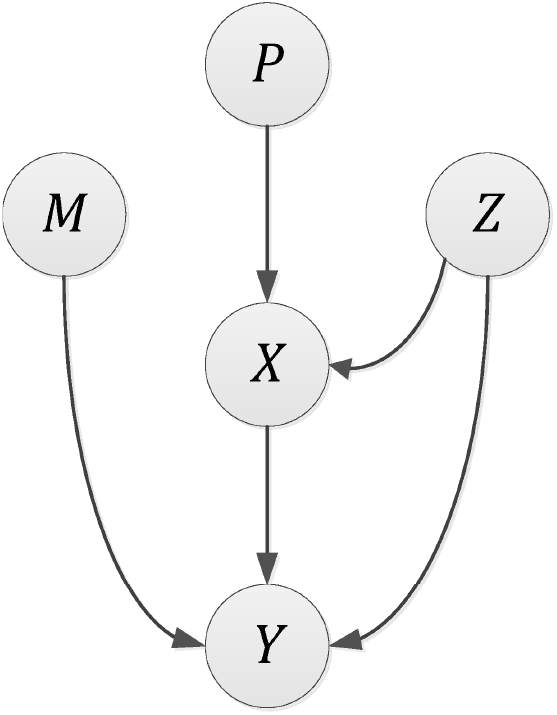
A Bayesian Network Model describing the relation between observed data: genotype matrix ***X***, disease infection labeles ***Y***, and unknown sub-population specific parameters: allele frequency matrix ***P***, membership information ***Z*** and hybrid disease model ***M***.

## 5 INFERENCE

In the Bayesian inference framework, we would like to obtain the posterior of the hidden parameters given the data, i.e. 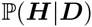. In this section, we present an algorithm based on the Markov-Chain Monte-Carlo (MCMC) method to achieve such a goal.

It is worth mentioning that the proposed statistical model presented in the preceding section can be viewed as a generalisation of the model used by STRUCTURE for unlabelled datasets. In particular, if we remove the labels from our model, the Bayesian inference amounts to unsupervised clustering of individuals based on their genotyped data which has been previously carried out in [9]. In a GWAS, however, we wish to incorporate labeled data samples and take advantage of the additional information provided by labels during the inference.

Conventional frameworks intend to correct for the effect of population stratification by first clustering the data samples, and then feeding each cluster of data into a GWAS module to infer causal factors in a separate phase. We show that clustering and finding causal disease factors are needed to be inferred simultaneously.

### 5.1 MAP Estimation via Gibbs Sampling

We set out to elaborate upon previously developed numerical methods to maximize 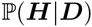, which indicates the posterior probability distribution of allele frequencies, subpopulation memberships and the disease models based on an observed dataset. In order to do so, we have taken advantage of the Markov-Chain Monte-Carlo (MCMC) methods, which not only have demonstrated top-notch performances in a variety of numerical optimization applications but are also easy to implement. These methods are extremely useful for obtaining samples from a probability distribution, especially in cases where the closed form formula for generating the samples is either unknown or too complex to be directly used, as in the case of our problem. In the following we briefly discuss an effective numerical technique in the MCMC family, known as *Gibbs Sampling*, which has been employed in this study.

There are a handful of problems in which a number of independent samples from a known high-dimensional distribution *π(θ_1_, θ_2_, θ_3_, …, θ_n_)* are needed. However, direct sampling from *π* is not numerically feasible. In such cases, Gibbs sampling guarantees generation of independent samples which converge to the desired distribution *π*, should the *ergodicity* condition is satisfied. The procedure for generating these samples is as follows:

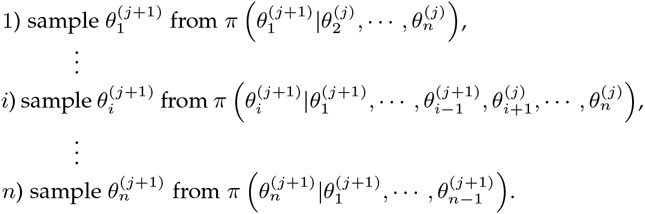

Performing steps 1 through *n* is called an iteration. It is shown that if the number of iterations required for convergence to the steady state, also known as the *burn-in* period, is sufficiently large then the Markov chain closely imitates the desired distribution. The number of iterations between two consecutive samples, shown by *c*, should also be sufficiently large. Fortunately, it is easy to show that these conditions hold for the problem at hand, thus making the MCMC method applicable to our algorithm.

The analogy between the model at hand and the Gibbs sampling framework mentioned above becomes clear by replacing (*θ_1_, θ_2_, θ_3_*) with (***P, M, Z***). However, our experimental observations confirm that maximizing the posterior distribution for disease models in the final stage of each iteration, instead of sampling from it, results in higher convergence rates. Hence, the sampling of the posterior probabilities can be done by iterating the following steps:

1. sample ***P***_(*m*+1)_ from 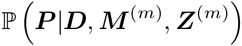,
2. sample ***Z***_(*m*+1)_ from 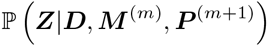,
3. find ***M***_(*m*+1)_ by maximizing 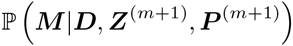,

where *m* denotes the index of previous iteration.

### 5.2 Inference of Allele Frequencies

Since minor allele frequencies ***P*** are independent of the disease model, the first step of the proposed inference algorithm can be simplified into sampling of ***P***^(*m*)^ from 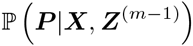. Recall from Section 4 that the prior distribution for allele frequencies, i.e. 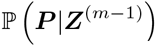, is modeled via a Dirichlet distribution with parameters *λ_1_, …, λ_J_*. Also, 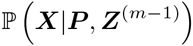 which resembles the probabilistic model for generating genotype data from MAFs is assumed to be a multinomial probability distribution. Hence, the posterior probability distribution for allele frequencies can be written as:

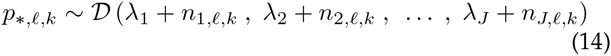

where 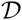 indicates a Dirichlet distribution with parameters *λ_j_* + *n_j,ℓ,k_*, *j* = 1, 2,ℓ, *J*. The relation in (14) directly follows from the fact that Dirichlet distribution is the conjugate prior for multinomial distribution. *λ*s are user-specific parameters while the quantities *n_j,ℓ,k_* are defined as:

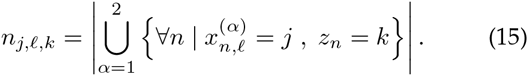

In other words, *n_j,ℓ,k_* represents the number of chromosomes in the *k*th sub-population of the dataset which contain the *j*th allele type in their *ℓ*th genetic locus. These parameters indicate the empirical abundance of specific allele types in each locus and sub-population.

### 5.3 Inference of Sub-population Memberships

In the second step of the algorithm, each person is assigned to a cluster based on current estimates of other target variables and also observed data. Mathematically speaking, one has to sample_)_from the posterior distribution 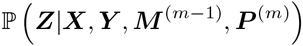 to obtain a modified estimate of the membership information ***Z***^(*m*)^.

In this stage we need to acquire a likelihood function for the disease label of each individual ***Y***, based on current estimates of disease model ***M*** and genotype data ***X***. That would resemble the use of *F_z_* (***X***) functions in our proposed model. In this step, 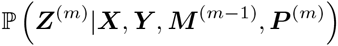 can be written as:

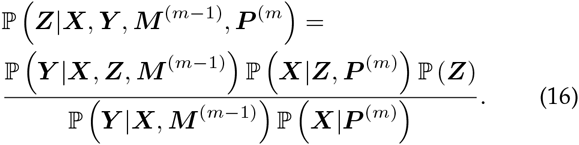

For the *n*th individual the equation reduces to the following form:

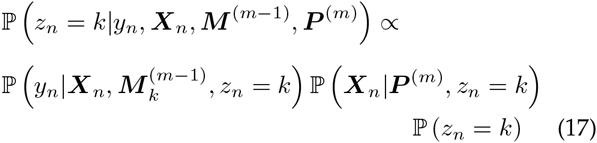

Once (17) is computed for all *k* ∈ {1,2,…,*K*}, we can normalize the quantities in order to attain a discrete probability distribution. 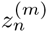 can then be achieved by sampling from this discrete probability distribution.

### 5.4 Inference of Disease Models

The final step of our modified Gibbs sampling procedure corresponds to finding the most probable disease models for each sub-population according to the posterior probability distribution of *M_k_s*, i.e. 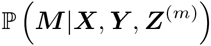. It should be reminded that given the genotype data of an individual, his/her infection to the disease is assumed to be independent from MAFs. A notable fact is that ***Mk*** may be inferred solely from {(***X***_*n*_, *y_n_*, *z_n_*) |*n* = 1, 2,…, *N*}. In this regard, the inference can be done by any of the previously introduced disease model identification methods in the GWAS literature. However, in this study we use our general disease model proposed in Section 4.2.

It is clear that the formulation 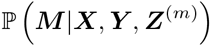 can be written as:

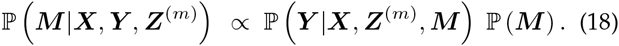

By replacing the equations from (13) into (18), one can alternatively have:

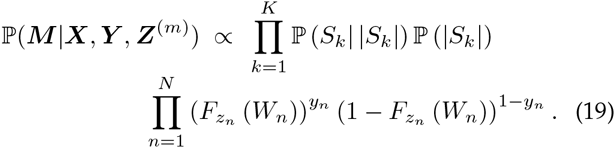

Disease model selection step implies the maximization of 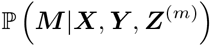 with respect to variables *S_k_* and *F_k_* (.) for all *k* = 1, 2, …, *K*. It is easy to investigate that maximization with respect to probabilities *F_k_* (.) has an analytical solution. Optimal disease probabilities at the *m*th iteration, 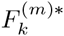, can be obtained via the following relation (calculations are presented in the Appendix A):

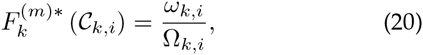

where 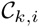 represents the *i*th combination of the causal factors in the *k*th sub-population. Obviously, for the *k*th sub-group we have 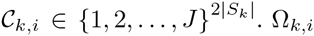 and *ω_k,i_* are defined as:

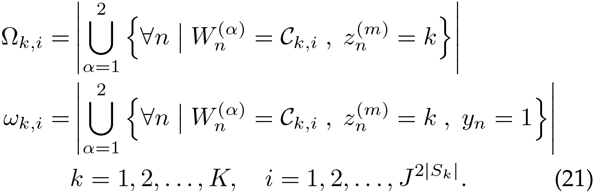

All 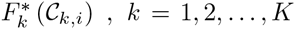 are calculated independently for each cluster as well as for each combination 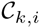. The intuition behind equation (21) seems obvious, since the probability of disease infection for a group of individuals with a particular allele combination and in a specific subpopulation is estimated by the empirical ratio of those who are infected, to the number of all the individuals having that combination. By substituting the optimal disease infection probabilities into (13), the following formulation is achieved:

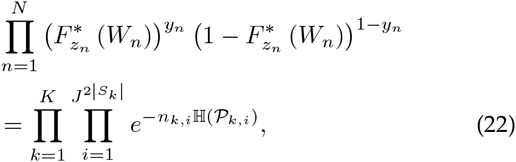

where *n_k,i_* denotes the number of chromosomes with 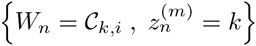, and 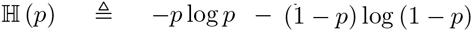 denotes the Shannon entropy. Likewise, 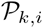 indicates the empirical ratio of disease infection in the *k*th sub-population for those individuals with the allele combination 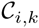. Again, the proof is given in Appendix A.

Maximization of (19) with respect to the remaining variables, i.e. the sets *S_k_*, *k* = 1, 2, …, *K*, does not have an analytical solution and should be determined via exhaustive searching in a valid solution space. This is an essential step in almost all GWAS methodologies. Mathematically speaking, for all possible choices of |*S_k_*| and loci in *S_k_* the following objective function should be evaluated and consequently maximized:

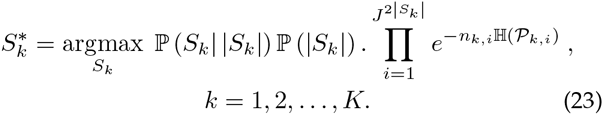

The above optimization problem, in its simplest form, requires a search on the total possible subsets of SNPs to be solved which renders this approach inapplicable even for moderate numbers of SNP loci. However, it should be noted that by choosing *η* (the expected number of genetic loci involved in the formation of disease) wisely, one can control the computational complexity of the search. In other words, genetic diseases are mostly caused by abnormalities in a limited number of SNPs, say < 10, rather then the whole set of SNPs in genome ~ 10^6^. This fact will significantly reduce the search space since for |*S_k_*| ≫ *η* the objective function in (23) becomes negligible and should not be checked. Moreover, by imposing the prior assumption regarding the consideration of epistasis only for neighboring SNPs (choosing relatively small values for epistasis length Δ) the valid search space will be reduced even further and the computational complexity of the optimization becomes practically tractable.

Under mild condition including the *ergodicity* criteria, one can investigate that the series

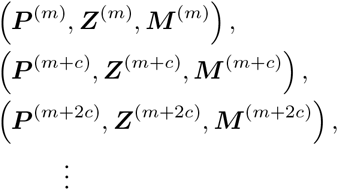

for sufficiently large *m* and *c* will resemble independent samples of the posterior distribution of the overall model. It should be noted that initial value of ***P***, ***M*** and ***Z***, denoted by ***P***^(0)^, ***M***^(0)^ and ***Z***^(0)^, are selected according to their corresponding prior probability distributions. Drawing ***Z*** from a uniform distribution seems reasonable unless some prior information including geographical, ethnic or racial characteristics of the individuals are present.

## 6 EXPERIMENTAL RESULTS

In this section, the experimental results of the proposed statistical framework are presented. Moreover, performance of our method has been compared with conventional GWAS methodologies as well as state-of-the-art clustering frameworks in the area of population genetics. We will show that the proposed framework surpasses both conventional GWAS algorithms and unsupervised clustering methods in determining the causal factors of the disease and also identifying the hidden population structures. The next part will discuss experimental results over synthetic data in addition to providing explanations regarding the generation of these datsets. Final part of the section is devoted to representation and analysis of computer simulations and comparisons.

### 6.1 Synthetic Data

In order to test the performance of our algorithm, we developed simulated data using our data generation model discussed in Section 4 whose hidden parameters were known prior to testing our framework. The data generation model takes into account realistic assumptions underlying living organisms such as population stratification, genetic barriers and linkage equilibrium.

For the sake of simplicity, we have assumed that the dataset consists of two hidden sub-populations, i.e. *K* = 2. Inspired by the attributes of real genotyped datasets, we have also assumed that most of the genotyped loci have same MAFs in both sub-populations, which are considered as random values in the range (0, 0.1). Consequently, only a small fraction, denoted by γ, of the loci have sub-population specific frequencies and thus can be useful during the clustering; However, these loci are not assumed to be known a priori. We have assumed *γ* = 5% in all of our simulations while the number of geotyped loci varies between 20 and 5000.

In the next phase, disease labels will be generated for each individual based on the statistical infection model discussed in the preceding sections. Causal factor numbers and corresponding genetic loci are determined through random sampling from prior distributions with *η* = 2 (expected number of causal loci) and Δ = 10 (the physical extent of linkage disequilibrium in genome). It is worth mentioning the causal loci are assumed to be different in each subpopulation. This assumption models the fact that several different malfunctions in the biological pathways lead to the same disease. Moreover, a number of possible allele combinations of causal factors are chosen to be disease causing, i.e. with 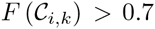 which implies a high risk of infection if 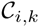 is exposed, while the other combinations are assumed to be neutral, i.e. 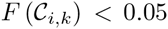. According to therapeutic properties of many complex diseases, combination of minor alleles at SNP loci with a moderate or higher linkage disequilibrium have been identified as the main causal factor of the illness[4], [5]. These assumptions are appropriately addressed during the data generation phase via parameter settings. Finally, it should be noted that the total number of iterations and the *burn-in* period for our MCMC implementation are set to 20000 and 10000, respectively.

### 6.2 Results

We have compared the performance of our method to STRUCTURE [9], in determining the hidden subpopulations within the dataset. STRUCTURE is known as the state-of-the-art unsupervised clustering algorithm for genotype data. The results are depicted in Fig. 2 and Fig. 3 for datasets of size *N* = 600 and 1000, respectively. STRUCTURE ignores the disease labels since its core algorithm is designated for unsupervised scenarios. However, we have observed that if use only the case group, i.e. the group with *y_n_* = 1, the performance of STRUCTURE will improve for large datasets. However, it is evident that for small number of loci, i.e. *L* < 1000, the proposed framework has a significantly improved performance over STRUCTURE and its variant. Moreover, in extreme scenarios STRUCTURE has an accuracy around 50% in a two-class problem which renders this method inapplicable in such cases. The mentioned supermacy for the proposed method is due to employment of disease labels and an appropriate disease model during the inference, while STRUCTURE only uses only allele types in informative loci.

**Fig. 2.**
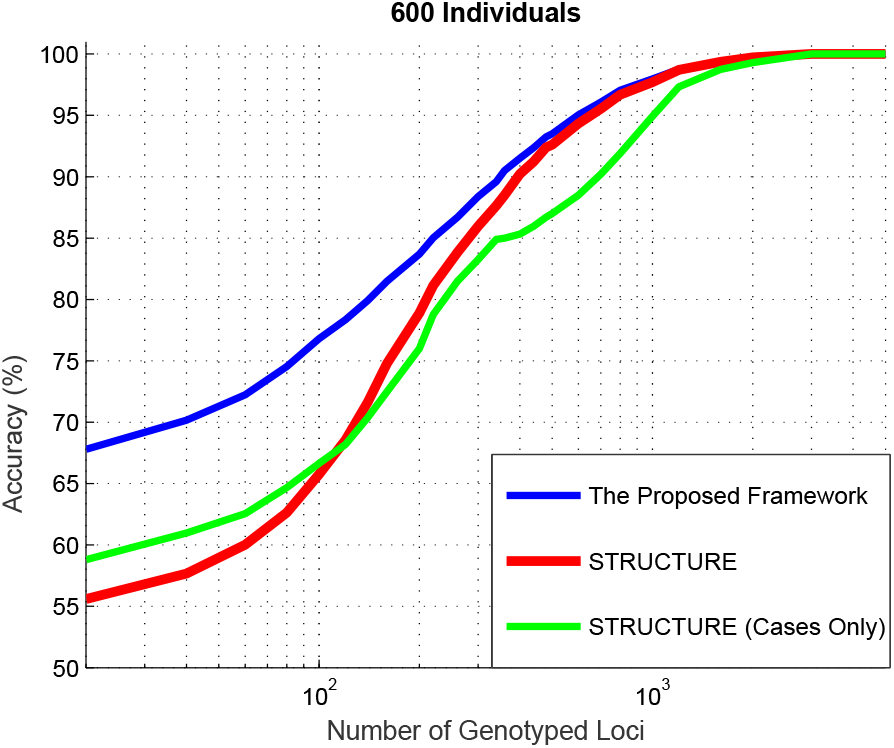
Accuracy in identification of hidden sub-populations as a function of the number of genetic loci, for our proposed method and the STRUCTURE framework with both all data points and cases only. Dataset consists of overall 600 individuals, and only 5% of genetic loci have different allele frequencies between the two hidden sub-populations. The proposed method surpassed the state-of-the-art, specifically in low loci number regimes.

**Fig. 3.**
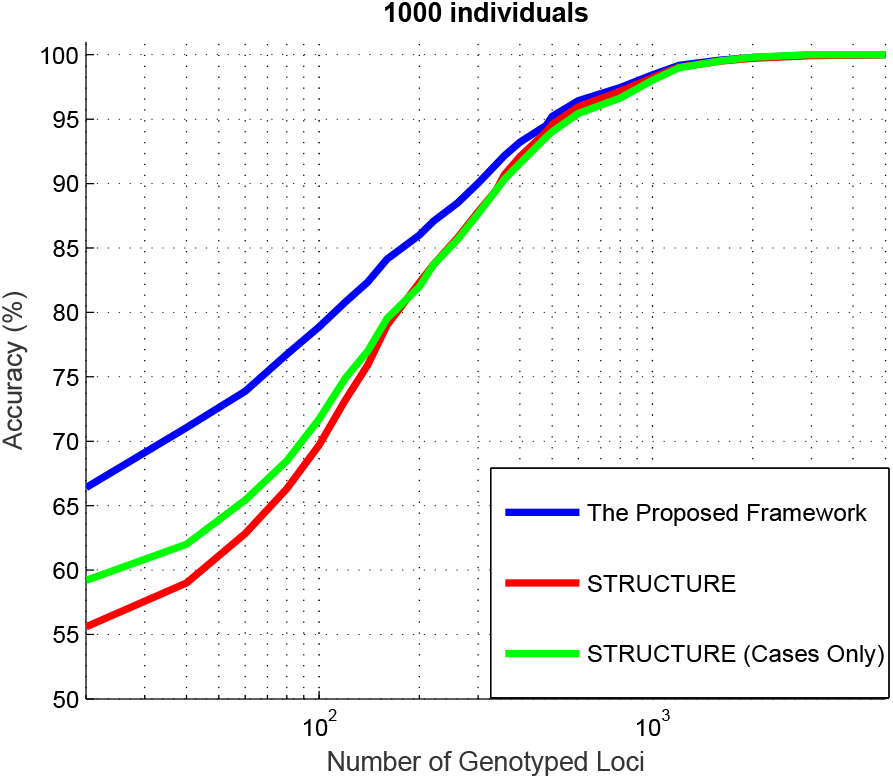
Accuracy in identification of hidden sub-populations as a function of the number of genetic loci, for our proposed method and the STRUCTURE framework with both all data points and cases only. Dataset consists of overall 1000 individuals, and only 5% of genetic loci have different allele frequencies between the two hidden sub-populations. As can be seen the performance of STRUCTURE in “cases only” mode has been improved, however, the proposed method still performs better.

Fig. 4 illustrates the region in *N-L* plane (number of individuals vs. number of genotyped loci) in which the methods have shown a clustering accuracy of 80% or higher. In this regard, the borders of this region is shown for the proposed method and STRUCTURE in blue and red colors, respectively. As it can be seen, the proposed method encompasses a relatively larger area in the plane which indicates the method outperforms unsupervised clustering algorithms when the number of individuals or the number of genotyped loci are small. It worth mentioning that for practical reasons it is common for researchers to reduce the number of genotyped loci in genome-wide association study since the inherent complexity of the problem usually scales exponential with *L*. On the other hand, the number of individuals in a GWAS dataset is limited due to financial issues in acquiring of the data samples.

**Fig. 4.**
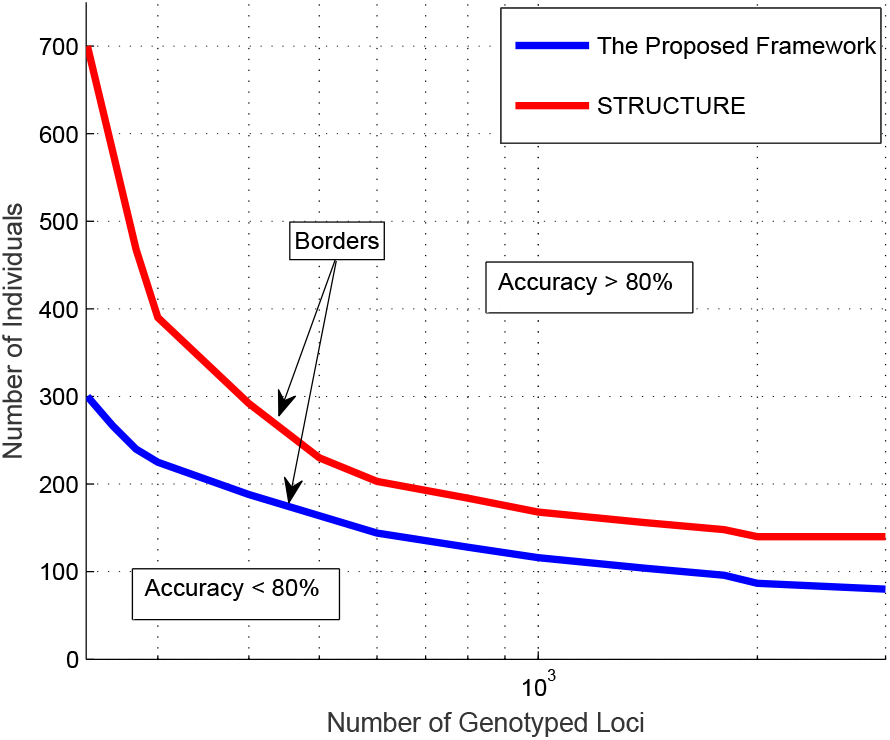
The area in *N-L* plane (number of individuals and number of genotyped loci) in which the accuracy of clustering methods are above 80% in a two-class problem. The proposed method is compared with the STRUCTURE framework. It can be seen that our method has an acceptable performance in a relatively larger area of the plane, implying a more robust performance for small size datasets. The achieved improvement is due to employing disease infection labels during the clustering.

An important aspect of any GWAS methodology is its capability for correct identification of disease-causing sites in a given dataset. The performance of the proposed framework is shown in Fig. 5 where a number of Manhattan plots are depicted to show the statistical significance level of the genetic loci being tested. By statistical significance we simply mean − log (p-value), where all p-values are computed according to the hypothesis of being a causal factor. As it can be seen, the significance level of the main causal loci corresponding to each of the hidden sub-populations are relatively small, hence, resulting into spurious inferences (blue plots). The main reason for this phenomenon is the lack of an appropriate population stratification to separate different case/control groups. As a result, different subpopulations would suppress the significance level of each other by introducing noisy signals. However, effective clustering of the dataset and computing the significance level of each causal factor only within its corresponding subpopulation achieves a considerably higher significance level and avoids mis-identifications. the latter can be achieved by our proposed framework while conventional clustering algorithms have been failed to effectively find the latent population structures.

**Fig. 5.**
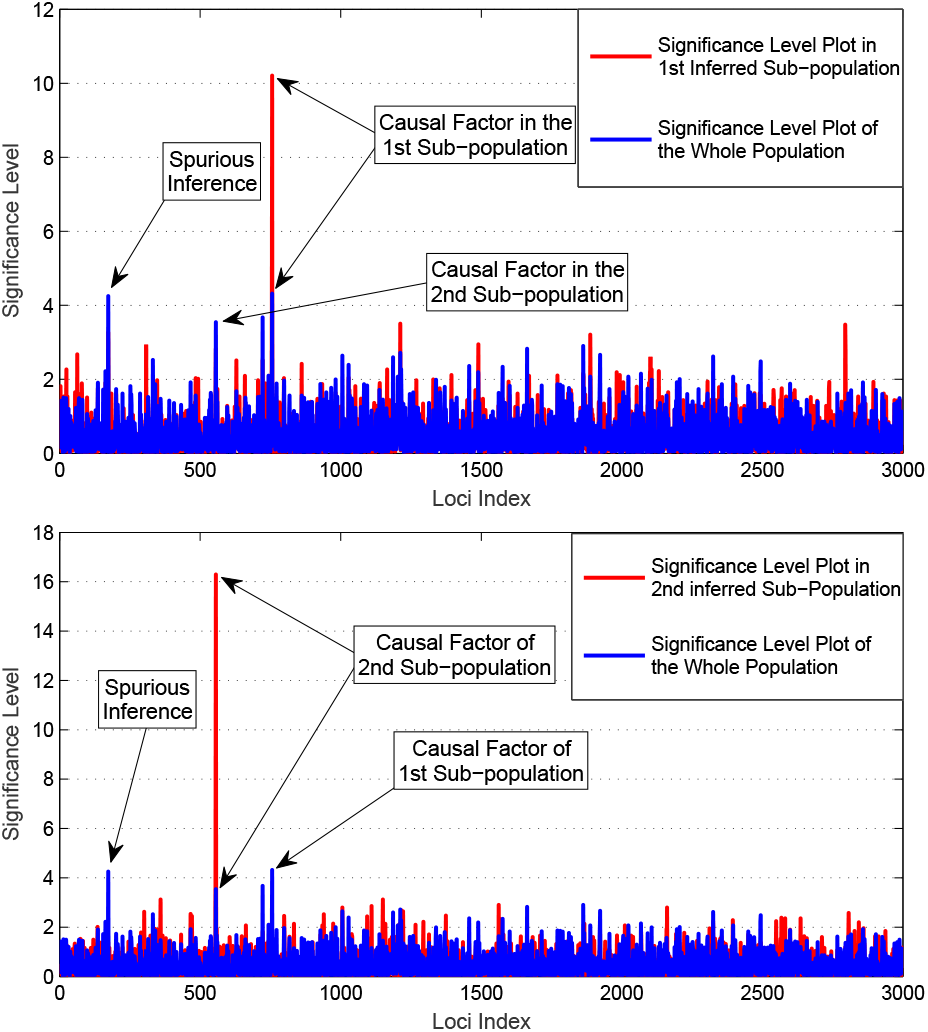
Comparison of disease-causing site identification via conventional GWAS (Blue) and the proposed clustering-based framework (Red). (Up): statistical significance level, i.e. − log (p-value), is computed over the whole population (Blue plot), and over the inferred 1st sub-population. (Down): the same procedure for the 2nd sub-population has been done. Evidently, conventional hypothesis testing without correction for the effect of population stratification results in small significance levels and spurious inferences. However, once the hidden sub-populations are correctly identified, disease-causing sites can be robustly inferred.

In order to further illustrate the performance of the proposed method we have compared its accuracy with five rival methodologies. The rival methods are simple single-locus and bi-locus hypothesis testing algorithms implemented in PLINK toolbox [55], an improved genetic algorithm (IGA) framework introduced in [56], GBOOST which is designated to capture mutual epistatic relations [57], simple hypothesis testing with a PCA-based population stratification module [41], and a GWAS method inspired by linear support vector machines (SVM) proposed in [7]. Two sets of datasets have been used for this experiment, which consist of *K* = 1 and *K* = 2 hidden sub-populations, respectively. The results have been shown in TABLE. 1. It should be noted that parameters for each method have been tuned to achieve the best performance. For the case of *K* = 1 almost all methods have an acceptable performance, while PLINK and GBOOST perform marginally better. However, it can be observed that when there are strong hidden population structures, as in the case of *K* = 2, the accuracy of the proposed framework has significantly surpassed the rivals. As a result, all the mentioned methodologies face with spurious statistical inferences which result in erroneous decisions. Also, it has been seen that PCA-based methods for population stratification are not useful in extreme cases.

**TABLE 1.** Performance comparison of the proposed framework with five rival methodologies in terms of accuracy in identification of causal factors. Two sets of datasets have been employed for this test which consists of *K* =1 and *K* = 2 hidden sub-populations, respectively. As can be seen, when *K* > 1 the proposed method outperforms rival algorithms.

|  | Accuracy |  |
| --- | --- | --- |
| | $K = 1$ | $K = 2$ |
| PLINK I | 64% | 19% |
| PLINK II | 93% | 27% |
| IGA | 32% | 7% |
| GBOOST | 94% | 30% |
| PCA-based | 65% | 34% |
| SVM-based | 71% | 13% |
| Proposed | 79% | 74% |

## 7 CONCLUSIONS

Population structures are shown to have a tremendous impact on the accuracy of many genome-wide association mapping studies. The majority of methods which are intended to rectify this shortcoming are based on unsupervised clustering of genotype data in a preprocessing stage to cancel the effect of population structures, and then feeding each cluster for a GWA study separately. This strategy confronts sever problems in small-size datasets since the MAFs do not necessarily suffice for robust identification of sub-populations. On the other hand, a variety of recent medical discoveries verify that most of complex diseases are multigene and thus may have several infection models according to genome. Based on this fact, this paper proposes a novel statistical framework to perform association mapping and population structure identification simultaneously and interactively. We have shown that in extreme scenarios, such as many real-world datasets, the accuracy of the proposed framework in identifying population structures can be improved up to 10% to 15%. Moreover, false discovery rate in association mapping stages are dropped dramatically.

In our future works, we will mainly focus on the effect of population admixtures, i.e. multiple genetic ancestries for each individual, which has been neglected in this study for simplicity. In addition, it is possible to transform the mathematical core of the current study from a parametric viewpoint, to a Bayesian non-parametric setting which is shown to be more robust against parameter configurations.

## Appendix A Analytic Solution for Disease Infection Probabilities

In this section we provide the proof for obtaining equations (21) and (22). In order to maximize 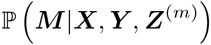, one can simply aim to maximize its logarithm:

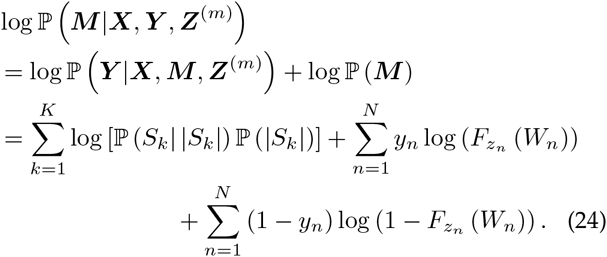

Obviously, only the two last summands depend on infection disease probabilities. Since *W_n_* is not continuous and takes only discrete values, one can rewrite the mentioned summands as:

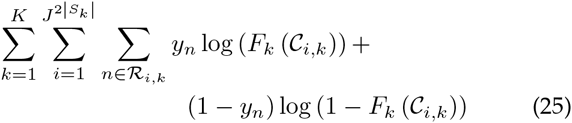

where 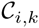 indicates the *i*th possible combinations of 2 |*S_k_*| alleles. The factor of 2 corresponds to the *diploid* assumption. Accordingly, 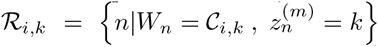. By calculating the derivatives of (25) with respect to 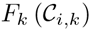 and omitting the constant factors, the above equation is simplified to:

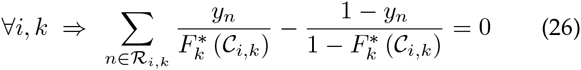

which alternatively means:

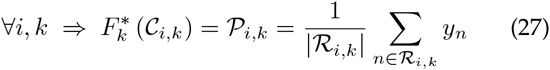

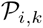 indicates the empirical disease infection ratio for individuals in 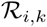. By substitution of the above optimal disease infection probabilities into (25), one simply achieves the following equation in terms of *S_k_* and the inputs of the original problem:

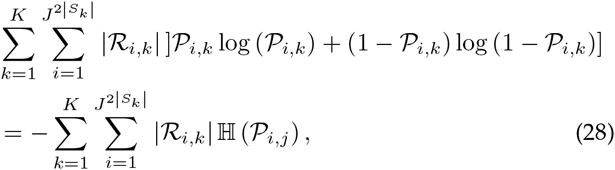

where 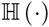 denotes the Shannon entropy. As a result, we have:

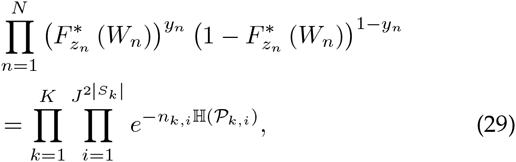

which is equation (22) and the proof is complete.

---

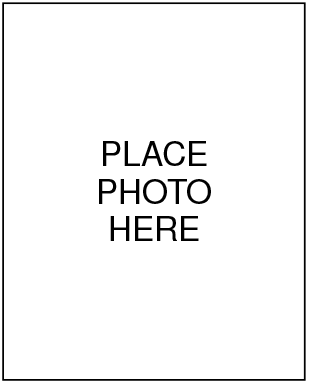
**Amir Najafi** received his B.Sc. and M.Sc. degrees from Electrical Engineering Dept. of Sharif University of Technology (SUT), Tehran, Iran, in 2012 and 2015, respectively. He is currently a Ph.D. student of Artificial Intelligence program at Computer Engineering Dept. of Sharif University of Technology. His research interests include bioinformatics, machine learning and information theory.

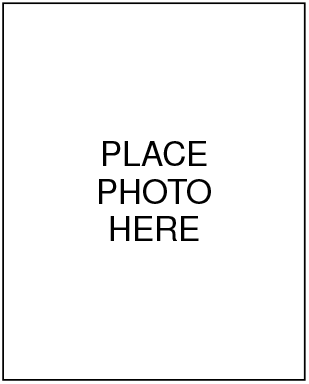
**Sepehr Janghorbani** received his B.Sc degree from Computer Engineering Dept. of Sharif University of Technology, Tehran, Iran, in 2016. He is currently a Ph.D. student in Rutgers University, NJ, USA. His research interests include neurosciences, medical imaging and bioinformatics.

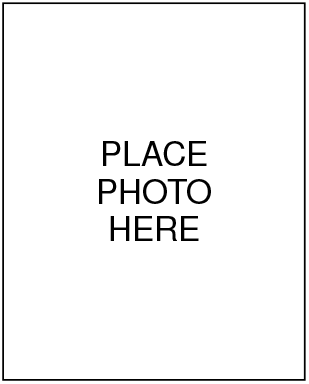
**Seyed Abolfazl Motahari** is an assistant professor at Computer Engineering Department of Sharif University of Technology. He received his B.Sc. degree from the Iran University of Science and Technology (IUST), Tehran, in 1999, the M.Sc. degree from Sharif University of Technology, Tehran, in 2001, and the Ph.D. degree from University of Waterloo, Waterloo, Canada, in 2009, all in electrical engineering. From August 2000 to August 2001, he was a Research Scientist with the Advanced Communication Science Research Laboratory, Iran Telecommunication Research Center (ITRC), Tehran. From October 2009 to September 2010, he was a Postdoctoral Fellow with the University of Waterloo, Waterloo. From September 2010 to July 2013, he was a Postdoctoral Fellow with the Department of Electrical Engineering and Computer Sciences, University of California at Berkeley. His research interests include multiuser information theory and Bioinformatics. He received several awards including Natural Science and Engineering Research Council of Canada (NSERC) Post-Doctoral Fellowship.

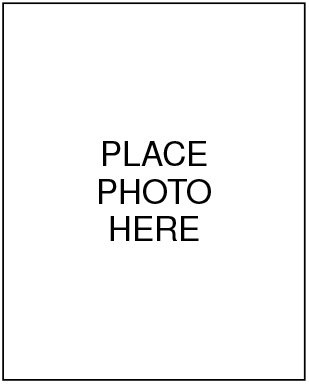
**Emad Fatemizadeh** is a faculty member of Biomedical Engineering in the Department of Electrical Engineering at Sharif University of Technology since 2004. He received a PhD degree from Tehran University. His research interests in biomedical engineering are in the areas of medical image analysis and processing, bioinformatics, statistical pattern recognition, medical data mining, and machine learning. Emad Fatemizadeh is a member of Institute of Electrical and Electronics Engineers (IEEE), board of founder of Iranian Society of Machine Vision and Image Processing (ISMVIP).

